# Hierarchical Value of Information in Microbial Predator-Prey Interactions

**DOI:** 10.64898/2026.08.26.747326

**Authors:** Peyman Fahimi, Michael Lynch

## Abstract

Information is fundamental to biological survival, but the amount of information and its biological value are not equivalent. Shannon information quantifies uncertainty reduction, whereas Volkenstein’s value of information measures how information changes the probability of a biologically relevant outcome. Although originally developed for molecular biology contexts, the latter concept has rarely been applied to environmental sensing and ecological interactions. Here we develop a value-of-information framework for microbial predator–prey interactions based on hydrodynamic sensing, in which prey detect fluid disturbances generated by approaching predators. Using a mechanistic model that incorporates sensory thresholds, memory, false alarms, biological benefits and costs, and predator encounter probability, we characterize mutual information from three hierarchical measures of biological value: encounter-conditional value, ecological value, and lifetime fitness value. The framework reveals how small amounts of sensory information can produce disproportionately large survival benefits during predator encounters, generating encounter-level value amplification in which biological value exceeds Shannon information. However, although global sensitivity analysis shows that such amplification is common, it is not universal and becomes progressively diluted at broader ecological and lifetime scales by encounter rarity, background noise, and sensory costs. Across most parameter combinations, the encounter-conditional value exceeded the ecological value, which in turn exceeded the lifetime fitness value. These results demonstrate that environmental sensing should be evaluated not only by how accurately it represents the external world, but by how strongly it changes biologically relevant outcomes. More broadly, the framework extends Volkenstein’s concept of information value to ecological interactions and provides a quantitative framework for predicting when environmental information enhances survival and fitness, thereby providing a platform for explaining the evolution, maintenance, diversification, and loss of sensory systems.

## Introduction

Living organisms continuously acquire information from their surroundings. Cells sense nutrients, toxins, light, temperature, viscosity, chemical gradients, mechanical cues, prey, predators, and conspecifics, and use this information to guide behavior and improve fitness (Levins 1968; Dusenbery 1992; Lynch 2024). For planktonic microorganisms, hydrodynamic disturbances generated by swimming predators can function as early-warning signals, allowing prey to initiate escape responses before capture (Buskey et al. 2002; Kiørboe 2011).

Classical information theory, introduced by Shannon, provides a powerful mathematical language for quantifying the amount of information transmitted through a communication channel (Shannon 1948). In biology, this framework has been widely applied to genetic sequences, biochemical signaling, sensory systems, and neural coding (Strait and Dewey 1996; Laughlin et al. 1998; Adami 2004; Gatenby et al. 2025; Gatenby 2026). In ecology, Shannon entropy has also been widely used as an index of species diversity (Pielou 1966; Peet 1974; He 2010; Pueyo 2007). However, although Shannon information quantifies uncertainty and also can be used as a measure of diversity, it is otherwise independent of specific meaning, utility, or biological consequence (Volkenstein 1983, 1994; Castanedo et al. 2024). Two signals may contain the same number of bits while having very different effects on survival of the recipient. Conversely, a weak or noisy cue may contain little Shannon information but still be extremely valuable if it is received at the right time and controls a high-stakes decision (Volkenstein 1983, 1994; Castanedo et al. 2024).

This distinction was emphasized by Mikhail Volkenstein, who argued that biology requires not only a theory of information quantity but also a theory of information value (Volkenstein 1983, 1994; Castanedo et al. 2024). Volkenstein defined the value of information in terms of the change in the probability of achieving a specified outcome after information is received (Castanedo et al. 2024). In this view, information value is not an intrinsic property of a message alone. It depends on the receiver, the context, the available actions, and the biological consequences of those actions. A signal is valuable when it changes the probability of survival, reproduction, successful development, or another biologically meaningful result.

Despite its conceptual importance, the value of biological information has remained largely undeveloped (Castanedo et al. 2024). Volkenstein applied the concept primarily to problems in molecular biology, including amino-acid sequences, mutations, and biopolymers (Volkenstein 1979, 1983, 1985, 1994). As a result, a central question of broader significance has remained underexplored: when does environmental information become biologically valuable?

Predator-prey interactions provide a natural setting for answering this question. For microorganisms living in fluid environments, predators generate hydrodynamic disturbances as they swim, and these disturbances can inadvertently function as early-warning signals for potential prey (Kiørboe and Visser 1999; Visser 2001; Lauga and Powers 2009; Kiørboe 2011). However, hydrodynamic sensing is not automatically beneficial. A prey organism must detect the predator early enough to escape, but excessive sensitivity may produce false alarms in noisy environments, while escape responses and sensory systems themselves carry costs (Dusenbery 1992; Laughlin et al. 1998; Dall et al. 2005; Niven and Laughlin 2008; Lynch 2024; Fahimi et al. 2026*b*). Moreover, sensory function can carry energetic costs even in the absence of an immediate stimulus; in adaptive sensory systems, maintaining the adapted state may require continuous energy dissipation (Lan et al. 2012). Thus, the value of hydrodynamic information emerges from a trade-off between encounter probability, early detection, warning time, false alarms, and the cost of response.

The biological systems envisioned here include planktonic protists and zooplankton. For example, ciliates can detect fluid deformation and initiate escape jumps from copepod predators (Jakobsen 2001), while copepods can detect hydrodynamic disturbances generated by approaching fish and escape before capture (Tuttle et al. 2019). The physical basis of such interactions builds on earlier theory describing encounter rates through sensory encounter distances (Gerritsen and Strickler 1977) and relating predator-generated hydrodynamic disturbances to mechanosensory perception thresholds (Kiørboe and Visser 1999; Visser 2001; Guasto et al. 2011).

Here, we develop a quantitative theory for the value of hydrodynamic information in microbial predator-prey interactions. Information content is quantified as the mutual information between predator presence and sensory response, whereas biological value is quantified by the improvement in survival produced by that response. We define three levels of value: the encounter-conditional value, which measures the survival benefit during an actual predator encounter; the ecological value, which weights this benefit by the probability of predator encounters; and the lifetime-fitness value, which compares the survival of information-using and non-information-using organisms. A hydrodynamic cue may therefore have a large encounter-conditional value because it enables escape from an imminent predator attack, while its ecological value is reduced when encounters are infrequent and its lifetime-fitness value reflects the long-term average survival advantage of information use. Thus, the biological value of the same sensory cue can differ substantially across biological scales. Under some conditions, relatively small amounts of Shannon information may generate disproportionately large biological consequences, a phenomenon we term value amplification.

Although the present study focuses on hydrodynamic predator avoidance in planktonic microorganisms, the underlying theoretical framework is considerably more general. Different prey organisms rely on diverse sensory modalities, including hydrodynamic, chemical, visual, mechanical, and electrical cues, depending on their biology and ecological context. The hydrodynamic example developed here serves as a representative case study, but any biological process in which organisms acquire environmental information to make threshold-based decisions—including prey detection, chemotaxis, nutrient acquisition, toxin avoidance, and other binary behavioral responses—can, in principle, be analyzed using the same value-of-information approach. We therefore view the present model as a first step toward a broader quantitative theory of the energetic and evolutionary value of environmental sensing across biological systems.

By extending Volkenstein’s concept from molecular biology to ecological sensing, this work provides a framework for asking not only how much information organisms acquire, but how much that information is worth. This shift is especially important for environmental sensing, where organisms do not need or cannot attain perfect representations of the world. They need actionable signals that improve survival under physical, ecological, and evolutionary constraints. By distinguishing information content from information value, our framework predicts when natural selection should favor investment in sensory systems, when simpler sensory strategies are sufficient, and when sensory information becomes evolutionarily disadvantageous because its benefits no longer outweigh its costs. For example, predation can favor increased investment in costly sensory structures, as illustrated by the evolution of larger eyes in *Daphnia* exposed to fish predation, presumably because enhanced predator detection compensates for the metabolic costs of greater sensory investment (Beston et al. 2019). Also, regressive evolution of vision in cave-dwelling animals provides a well-known example in which the maintenance and developmental costs of eyes are no longer offset by fitness benefits in permanently dark environments (Jeffery 2009). However, not all mechanosensory capabilities require dedicated sensory structures. In bacteria, the flagellum functions not only as a propulsive organelle but also as a mechanosensor that detects changes in mechanical load associated with surface contact and fluid resistance, illustrating how environmental information can be acquired using structures that already serve another primary function (Belas 2014). Thus, the selective maintenance or loss of mechanosensing depends not only on the balance between its costs and benefits, but also on whether mechanosensory function requires specialized investment or can be integrated into multifunctional biological structures.

Because sensory systems require resources for their construction, maintenance, and operation, their biological benefits must be evaluated together with their costs. Representative empirical estimates of the energetic costs of sensing across biological systems are summarized in Box 1 and motivate the sensory-cost term introduced in the net fitness-payoff model below. In addition to the energetic costs of sensing discussed in Box 1, the prey net fitness-payoff function developed in this paper includes several other costs and benefits, including the energetic cost of escape responses, the costs associated with stale information and false alarms generated by background noise, and the survival benefits of successfully detecting and escaping predators, all of which are described in detail in the next section. Because our objective is to develop a general theoretical framework for microbial plankton predator– prey interactions rather than a model for any particular species, we use representative values for these costs and benefits. Specifically, we explore broad ranges for all model parameters by systematically scanning the parameter space to encompass scenarios in which the costs of sensing outweigh its benefits, as well as those in which the benefits exceed the costs (Table 1—Appendix). This general framework can readily be adapted to species-specific systems by replacing the representative parameter values with empirically measured estimates.

The overall conceptual structure of the framework is illustrated in Figure 1. Predator-generated hydrodynamic disturbances are integrated by a sensory system possessing memory and a detection threshold, producing a binary sensory response. This response determines escape success and survival probability, from which biological value is quantified at encounter, ecological, and lifetime scales. Comparison of these value measures with Shannon mutual information allows the identification of value-amplification regimes in which biological value exceeds informational content.

**Figure 1:**
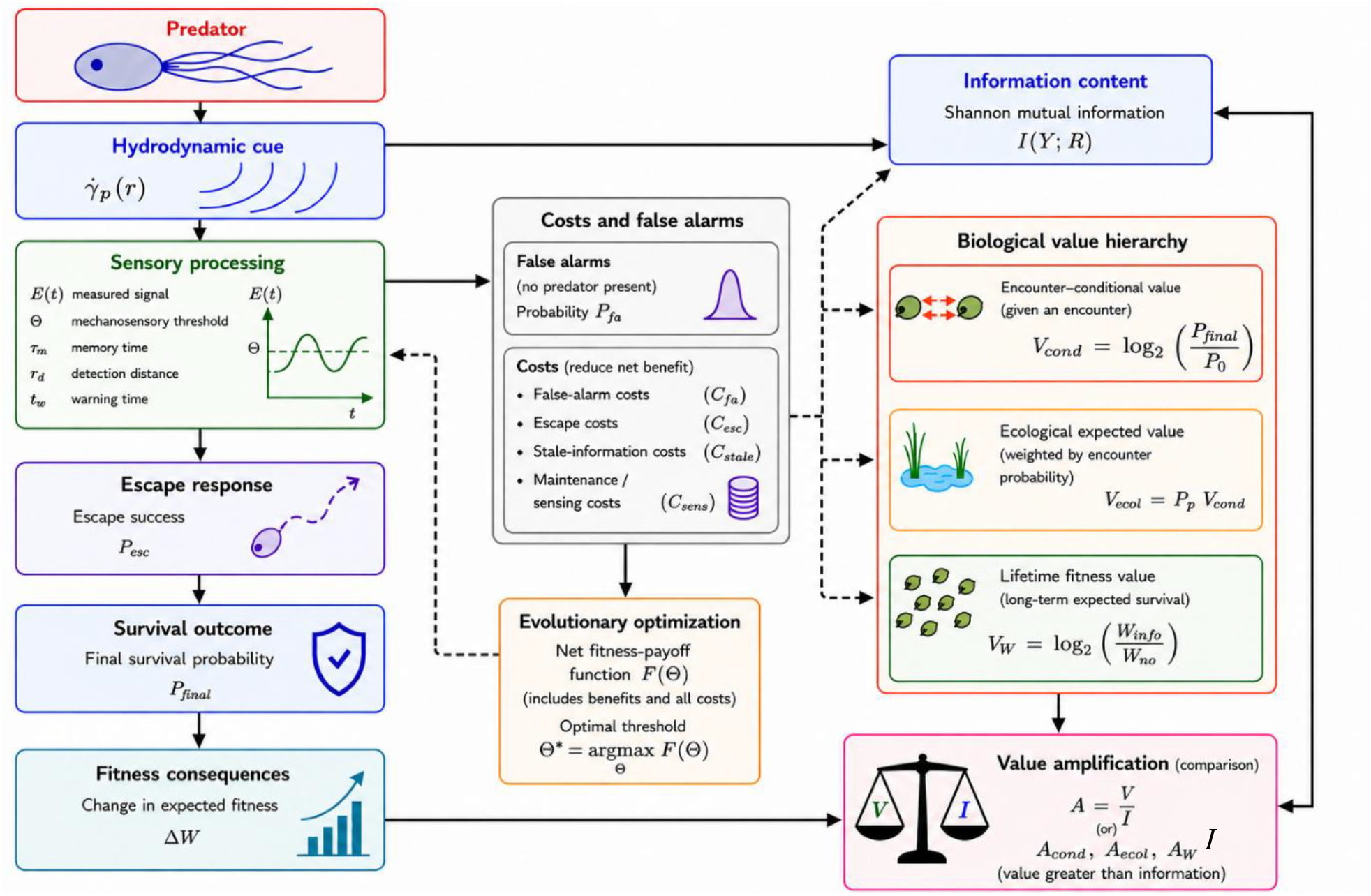
Conceptual framework for the hierarchical value of hydrodynamic information in microbial predator-prey interactions. Predator-generated hydrodynamic disturbances are sensed and integrated through a memory-dependent mechanosensory system characterized by an evidence variable and detection threshold. Sensory processing produces a binary detection response that influences escape success and ultimately survival probability. Evolutionary optimization determines the mechanosensory threshold by maximizing a net fitness-payoff function that incorporates the benefits of successful predator avoidance together with the energetic costs of sensing, false alarms, escape, and stale information. The resulting evolutionarily optimal threshold is subsequently used to evaluate both the Shannon mutual information and the biological value of information. The sensory response, together with predator presence or absence, defines the Shannon mutual information between the environmental state (Y) and the sensory response (R). Biological value is quantified at three nested scales: encounter-conditional value (V_cond_), which measures the survival benefit during a predator encounter; ecological expected value (V_ecol_), which weights this benefit by the probability of predator encounters; and lifetime fitness value (V_W_), which quantifies the long-term average survival benefit across repeated predator and non-predator periods. Comparison of these value measures with Shannon mutual information yields value amplification factors that quantify situations in which biological value exceeds informational content.

### BOX 1. Energetic Costs of Environmental Sensing

Environmental sensing imposes metabolic costs associated with both the construction and operation of sensory and signal-transduction systems (Lynch 2024). Construction requires the synthesis of receptors, signaling proteins, kinases, phosphatases, and other molecular components. For one representative mammalian phosphorylation-based signal-transduction system, the synthesis of its kinase and phosphatase components has been estimated to require approximately 2 × 10^11^ ATP molecules (excluding the interconvertible enzyme) (Lynch 2024), corresponding to about 0.2% of the total ATP investment required to construct a typical 5000 μm^3^ mammalian cell based on the cell-size scaling relationship of (Lynch and Marinov 2017). During operation, sensing consumes energy through the continuous ATP-dependent phosphorylation and dephosphorylation of signaling proteins required to maintain signaling activity, while some systems incur additional costs through processes such as receptor methylation and demethylation in bacterial chemotaxis. For the representative signal-transduction system of a 5000 μm^3^ mammalian cell described above, ATP consumption associated with kinase activity has been estimated to range from approximately 10^11^ – 12^12^ ATP molecules per cell cycle, representing up to 5% of the basal cellular maintenance energy budget (Lynch 2024).

Laughlin et al. (1998) investigate visual sensing in the blowfly retina, focusing on photoreceptors (R1-6), large monopolar cells (LMCs), and the photoreceptor-LMC chemical synapse. The authors show that the energetic cost of sensory processing is dominated by maintaining ion gradients via Na^+^/K^+^ pumps that support electrical signaling. Additional energetic costs arise from the phototransduction second-messenger cascade (including phosphorylation reactions and Ca^2+^ transport), chloride pumping in LMCs, and synaptic processes including neurotransmitter uptake, vesicle refilling, presynaptic Ca^2+^ pumping, and vesicle recycling. A single photoreceptor is estimated to consume 7.5 × 10^9^ ATP molecules s^−1^. To express this energetic cost per unit of information transmitted, the information transmission rate, *I* (bits s^−1^), is calculated from Shannon’s information-rate equation (Shannon 1949; Laughlin et al. 1998),

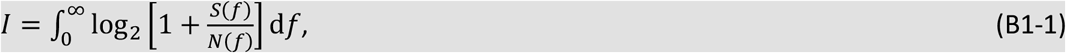

where *S*(*f*) and *N*(*f*) are the signal and noise power spectra at frequency *f*, respectively. The origin of this expression can be understood by treating each narrow frequency interval as an independent noisy communication channel. At a given frequency, the total received power is *S*(*f*) + *N*(*f*), whereas *N*(*f*) represents the uncertainty due to noise alone; their ratio is therefore 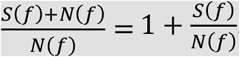. Shannon showed that the information carried by such a Gaussian channel increases logarithmically with this ratio, giving the term 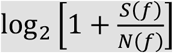. Because different frequency intervals contribute information independently, their contributions are summed over the entire frequency spectrum, which in the continuous limit gives the integral in Equation (B1-1) (Shannon 1949). Dividing the ATP consumption rate by *I* yields the metabolic cost per bit. For the photoreceptor, *I* ≈ 10^3^ bits s^−1^, corresponding to approximately 7 × 10^6^ ATP molecules per bit. Similarly, an LMC consumes 1.4–4.1 × 10^9^ ATP molecules s^−1^, corresponding to 9 × 10^5^–3 × 10^6^ ATP per bit, whereas a chemical synapse requires 2 × 10^4^–6 × 10^4^ ATP per bit. Remarkably, the authors estimate that photoreceptors alone consume about 8% of the total resting metabolic rate of the fly, and when the LMCs are included, visual information processing approaches 10% of the animal’s total resting energy budget, highlighting that sensory information processing constitutes a substantial fraction of whole-organism metabolism.

MacIver et al. (2010) quantitatively analyze the black ghost knifefish and its whole-body electrosensory system during prey search. They identify several sources of energetic cost associated with sensing: maintaining metabolically expensive neural and sensory tissue, generating the electric field required for active electrolocation, adopting a drag-inducing swimming posture that improves sensory coverage, and the neural processing needed for movable sensory organs. The largest reported sensory cost is the generation of the electric field, which consumes 3–22% of the fish’s total metabolic rate (up to ∼80 J/day for a fish with a 350 J/day energy budget), making active electrosensation a substantial metabolic investment.

Govern and ten Wolde (2014) analyze the chemotaxis signaling system of *E. coli* as a model of cellular sensing. They identify two energetic costs associated with sensing: (1) constructing the sensing machinery by synthesizing chemotaxis proteins, requiring approximately 2–8 × 10^7^ ATP per cell cycle, and (2) operating the sensing machinery through continuous ATP-driven phosphorylation/dephosphorylation reactions in the CheA–CheY signaling pathway, consuming approximately 1–5 × 10^7^ ATP per cell cycle (about 2,000–15,000 ATP s^−1^). For an *E. coli* cell with a volume of 1 μm^3^ and a one-hour cell cycle, the Lynch–Marinov scaling relationships (Lynch and Marinov 2017) predict a whole-cell construction cost of approximately 2.7 × 10^10^ ATP molecules and a maintenance cost of approximately 3.9 × 10^8^ ATP molecules per cell cycle. Thus, constructing the chemotaxis machinery represents approximately 0.07%–0.30% of the total cellular construction cost, whereas operating it accounts for approximately 2.6%–12.8% of the cellular maintenance energy budget. The pathway’s operating and construction costs are therefore of the same order of magnitude, although their relative contributions to the corresponding whole-cell energy budgets differ substantially.

Lan et al. (2012) analyze the energetic cost of sensory adaptation, primarily in the *E. coli* chemosensory system, while discussing yeast osmotic sensing and mammalian olfactory and visual adaptation as additional examples. They demonstrate that maintaining an adapted sensory state is an inherently non-equilibrium process requiring continuous energy dissipation. Quantitatively, for an *E. coli* cell containing approximately 10^4^ chemoreceptors, the authors estimate that maintaining the adapted state of the chemotaxis receptor system requires approximately 10^3^ ATP molecules per second; this estimate applies specifically to chemoreceptor adaptation rather than to the cell’s complete sensory machinery. The energetic maintenance cost of adaptation is estimated to be 5– 10% of the energy required to power a single flagellar motor rotating at 100 Hz.

### Theory

#### Value of hydrodynamic information

The concept of information permeates modern biology (Avery 2021). Terms such as genetic code, genetic information, transcription, translation, and cellular signalling all describe the storage, transmission, and processing of information by living systems. DNA can be viewed as a repository of biological information that is transmitted through RNA to proteins (Koonin 2011), while sensory systems continuously acquire information from the external environment to guide behaviour and improve fitness (Dusenbery 1992). Consequently, information theory, originally developed in engineering (Shannon 1948), has found widespread application in biology.

Our framework combines two complementary descriptions of information: Shannon information, which quantifies uncertainty reduction, and Volkenstein’s information value, which quantifies the effect of information on the probability of a biologically relevant outcome. Their conceptual distinction is summarized in Box 2.

##### BOX 2. Shannon Information and Biological Information Value

The mathematical foundations of modern information theory were established by Claude Shannon in 1948 (Shannon 1948). Shannon sought to answer a purely engineering question: how much information can be transmitted through a noisy communication channel? In Shannon’s framework, information is interpreted as a measure of uncertainty associated with the outcome of a random event. Rather than postulating a mathematical expression for this quantity, Shannon began by requiring that any measure of uncertainty satisfy three intuitive properties: (i) it should be continuous in the event probabilities; (ii) if all outcomes are equally likely (*p*_*i*_ = 1/*n*), it should increase monotonically with the number of possible outcomes *n*; and (iii) if a choice is decomposed into two successive choices, it should equal the probability-weighted sum of the uncertainties associated with the successive choices (Shannon 1948). Shannon proved in Appendix 2 of his seminal paper that these axioms uniquely determine the uncertainty measure (up to a positive multiplicative constant) to be the logarithmic form (Shannon 1948),

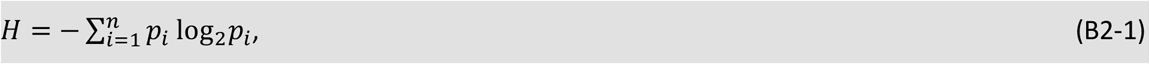

where *p*_*i*_ denotes the probability of observing the *i*-th possible outcome. Intuitively, an outcome with probability *p*_*i*_ carries −log_2_*p*_*i*_ bits of information: independent probabilities multiply, while their information should add, which leads naturally to the logarithm. Shannon entropy is then the probability-weighted average of this information over all possible outcomes. The logarithm is conventionally taken in base two so that information is measured in bits (binary digits), although other logarithmic bases merely change the units (for example, natural logarithms yield nats). Shannon entropy is maximal when all possible outcomes are equally probable and decreases as the probability distribution becomes more predictable.

Shannon’s theory provided a rigorous framework for quantifying the information content of DNA, RNA, and proteins (Johnson 1970; Strait and Dewey 1996). The frequencies of nucleotides or amino acids, sequence redundancy, and coding efficiency can all be analyzed using variants of Shannon entropy, although the precise meaning can vary with context. Shannon’s theory measures how much uncertainty a message removes, without considering what the message means or what effect it has. Thus, two messages containing the same number of bits are equivalent in Shannon’s framework even if one has major biological consequences and the other has none.

This limitation motivated Mikhail Volkenstein to introduce the concept of the value of biological information (see Eq. (1) in the main text) (Volkenstein 1994). He illustrated the distinction between information content and its value using the example of a traffic light: one bit of colour information carries the same Shannon information whether it controls traffic on a quiet side street or a busy avenue, yet its practical consequences are dramatically different (Makarov et al. 1987). Thus, identical quantities of Shannon information may possess vastly different biological values depending on the receiving system and its circumstances.

Volkenstein proposed measuring the value of information by its effect on the probability of achieving a specified biological outcome. Because the value of information depends on the objective being considered, the desired biological outcome must first be specified. If *P*_0_ and *P*_final_ denote the probabilities of achieving that outcome before and after the receipt of information, respectively, then the biological value of information is defined as (Volkenstein 1983, 1994; Castanedo et al. 2024)

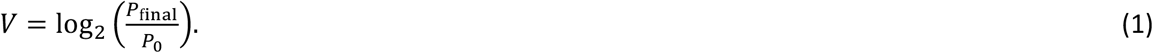

This form measures the proportional, rather than absolute, change in the probability of the desired outcome. The logarithm makes successive multiplicative changes additive, while base 2 places the value on the same logarithmic scale as Shannon information. Because *P*_final_ ≤ 1, the maximum possible value is 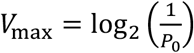; thus, the potential value of information decreases as the baseline probability *P*_0_ increases. Positive values of *V* indicate that the received information increases the probability of achieving the specified biological outcome, negative values indicate that it decreases this probability (for example, because the information is misleading), and zero indicates that the information has no effect on the probability of achieving the outcome. The specified outcome may range from complex processes, such as successful reproduction or predator avoidance, to much simpler cellular objectives, including ion or nutrient uptake, movement up a chemical gradient during chemotaxis, or maintenance of cellular homeostasis.

In the present work, the biological objective will be viewed through the lens of survival during a predator encounter. Accordingly, *P*_0_ corresponds to the probability of surviving without exploiting mechanosensory information, whereas *P*_final_ is the probability of survival after sensory information has been acquired and acted upon. Equation (1) therefore represents a direct application of Volkenstein’s general theory to predator-prey interactions and forms the foundation for our subsequent extensions to ecological and lifetime fitness measures of the value of information.

#### Hydrodynamic signal generated by a predator

Following (Visser 2001), when the predator–prey separation exceeds several predator body lengths (the far field), the fluid motion experienced by the prey due to the swimming predator can be approximated by a stresslet. In simple terms, a swimming organism pushes and pulls on the surrounding water as it propels itself, creating a disturbance that spreads outward and weakens with distance. Because the forces associated with propulsion and resistance balance one another for a freely swimming organism, the leading contribution to this distant flow is a stresslet. In this approximation, the velocity disturbance decays as 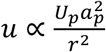, where *U*_p_ (m s^−1^) is the predator swimming speed, *a*_*p*_ (m) is a characteristic predator linear dimension (e.g., body radius or length), and *r* (m) is the radial distance from the origin of the hydrodynamic disturbance (approximated by the predator’s center) to the prey’s mechanoreceptor. Because mechanoreceptors respond to fluid deformation rather than fluid velocity itself, the relevant hydrodynamic signal is the local deformation (strain) rate, which scales with the spatial gradient of the velocity field. Since the stresslet velocity decays as *u* ∝ *r*^−2^, taking its spatial derivative gives a deformation rate that decays one power faster, 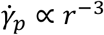. Here, 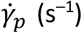 represents the intensity of the hydrodynamic cue experienced by the prey: larger values correspond to stronger fluid deformation and hence stronger stimulation of the mechanosensory system. We therefore write

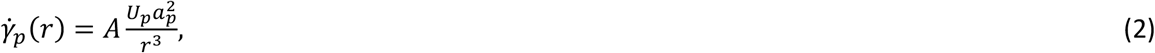

where *A* is a dimensionless geometric coefficient that accounts for the specific flow geometry and orientation of the predator-prey interaction. The stresslet description is a far-field approximation and therefore provides an idealized representation of predator-generated flows. At shorter predator-prey separations, higher-order hydrodynamic structures and organism-specific flow geometries may modify the deformation-rate field. The present formulation is intended to capture the dominant scaling of hydrodynamic signal strength with predator size, speed, and separation distance rather than to resolve detailed near-field flow structure.

#### Sensory memory and signal integration

Environmental sensing is fundamentally an information-acquisition process in which organisms continuously sample their surroundings to guide behavioral and physiological decisions. Regardless of whether the environmental cue is chemical, mechanical, or physical, sensory systems must balance three competing objectives: sensitivity, accuracy, and response speed (Lynch 2024). Because environmental signals are inherently noisy, reliable perception generally requires integrating sensory information over finite time intervals rather than responding instantaneously to individual receptor events. This principle has been developed extensively for chemosensing, where cells estimate external ligand concentrations by temporally averaging stochastic receptor occupancy (Berg and Purcell 1977; Endres and Wingreen 2008, 2009; Lynch 2024). Although our focus is mechanosensing rather than chemosensing, the same theoretical principle applies: prey estimate the presence of an approaching predator by integrating noisy hydrodynamic deformation signals over time. We therefore represent sensory memory using a finite-time integration of the incoming mechanical cue.

Temporal integration is a common feature of biological sensing systems, in which sensory signals are accumulated over finite time while previously acquired information gradually fades, providing a form of sensory memory. For example, bacterial chemotaxis can be described using a dynamical memory variable representing receptor methylation, whose temporal evolution governs adaptation to changing environmental signals (Tu et al. 2008). More generally, sensory and decision-making systems are commonly represented by leaky integrators that accumulate incoming signals while continuously forgetting older ones (Usher and McClelland 2001). The optimal balance between memory and forgetting is expected to depend on the statistical structure of environmental cues and the costs and benefits of retaining past information, and therefore may differ among organisms and ecological contexts. Because the physiological mechanisms underlying temporal integration in planktonic mechanoreceptors remain unknown, we adopt the simplest phenomenological model that captures these two processes. Specifically, we introduce a dimensionless internal state variable, *E*, representing the accumulated hydrodynamic evidence, whose dynamics are described by

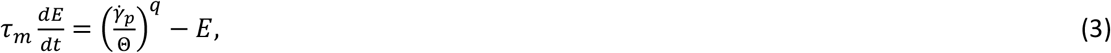

where *τ*_*m*_ (s) is the characteristic sensory memory timescale, Θ (s^−1^) is the mechanosensory threshold, defined as the deformation rate required to elicit a reliable sensory response, and *q* characterizes the nonlinearity of sensory transduction. Values *q* = 1, *q* > 1, and *q* < 1 correspond to linear, cooperative, and compressive sensory responses, respectively. The quantity constituting accumulated sensory evidence depends on the sensing mechanism. In chemosensing models, for example, environmental information can be accumulated over a finite observation period through receptor statistics such as the number of ligand-binding events, receptor occupancy over time, or the durations of unbound intervals between binding events (Endres and Wingreen 2009; Mora and Wingreen 2010). Here, *E* plays an analogous phenomenological role by integrating the hydrodynamic deformation signal over time.

The deformation rate 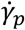 is normalized by the sensory threshold Θ so that the input is dimensionless and represents the stimulus intensity relative to the organism’s sensory capability. Consequently, the normalized input equals unity when the deformation rate is exactly at the detection threshold. Dividing both sides of Eq. (3) by *τ*_*m*_ shows that the first term, 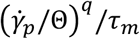, represents the normalized sensory input driving the rate of evidence accumulation, whereas the second term, *E*/*τ*_*m*_, represents the passive rate of evidence decay. Within this phenomenological framework, the decay is assumed to be first order. Consequently, in the absence of stimulation 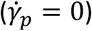, the internal state *E* relaxes exponentially toward zero with characteristic timescale *τ*_*m*_. The plausibility of such exponential decay of a sensory-memory variable is supported by bacterial chemotaxis: in *E. coli*, receptor methylation acts as a memory variable, and deviations from the adapted methylation level decay exponentially near steady state with an intrinsic adaptation timescale (Tu et al. 2008). Equation (3) constitutes the primary sensory model used throughout this study, and unless otherwise stated, all detection probabilities are obtained by numerically solving this dynamical equation.

#### Analytical approximation for detection range

For analytical tractability (as shown in the workflow in Fig. 2), we first consider the quasi-steady limit in which sensory integration occurs much faster than changes in predator-prey separation. In this limit, the internal sensory variable rapidly relaxes to its steady-state value,

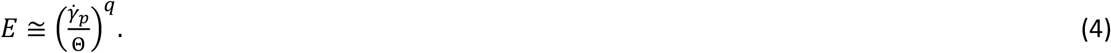

**Figure 2:**
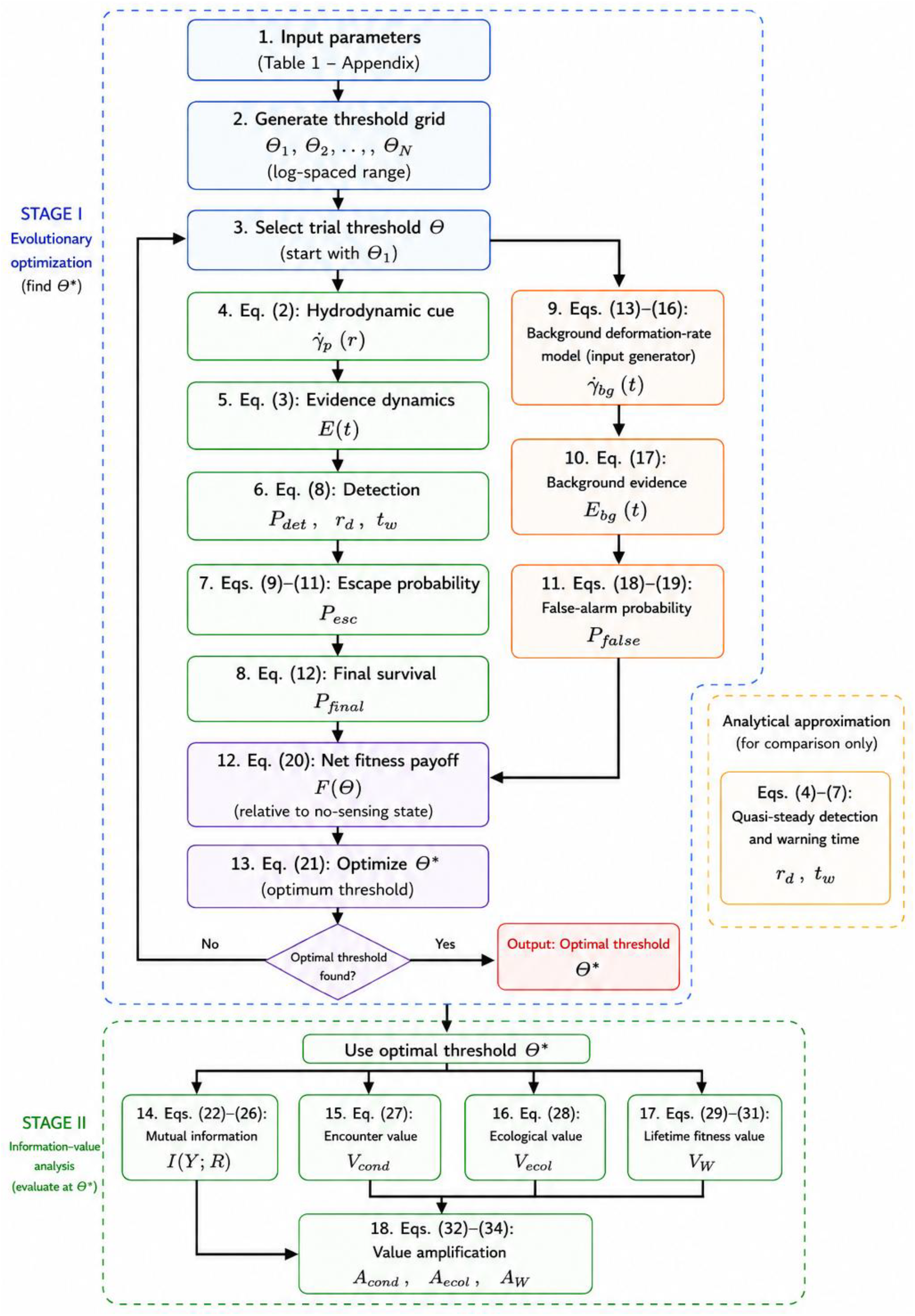
Computational workflow of the proposed framework. Stage I identifies the evolutionarily optimal mechanosensory threshold by maximizing the net fitness-payoff function, whereas Stage II evaluates the corresponding information-theoretic and biological value measures using the optimized threshold. The analytical approximation (Eqs. 4–7) is used only for comparison and is not part of the optimization procedure.

Because the sensory input in Eq. (3) is normalized by the mechanosensory threshold Θ, the accumulated evidence *E* is dimensionless and measured relative to the organism’s detection threshold. Consequently, *E* = 1 corresponds to the accumulation of sufficient normalized evidence to reach this threshold. We therefore assume that prey detects the predator once the accumulated evidence reaches or exceeds unity. Evaluating the quasi-steady relation in Eq. (4) at the detection distance *r* = *r*_*d*_ and setting *E* = 1 gives 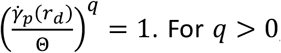. For *q* > 0, taking the *q*^*th*^ root of both sides gives

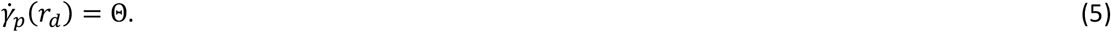

In the quasi-steady limit, the sensory nonlinearity *q* affects the rate of evidence accumulation but does not alter the detection threshold itself. Substituting Eq. (2) into Eq. (5) yields

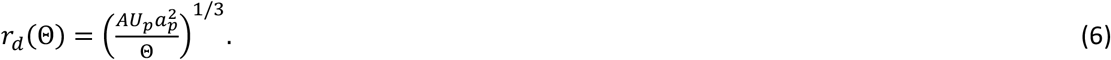

Eq. (6) shows that detection distance depends only weakly on predator swimming speed and sensory threshold, scaling as 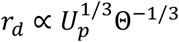, but more strongly on predator size, as 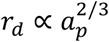. For example, an eightfold decrease in the mechanosensory threshold doubles the detection distance, whereas doubling predator size increases it by about 59%. This equation yields a closed-form expression for the detection range and provides useful biological intuition regarding the effects of predator size, predator speed, and sensory threshold. Equations (4)–(7) provide analytical approximations that are used solely for comparison with the full dynamical solution described in the framework (Figure 2) and evaluated in the results (Figure 3).

**Figure 3:**
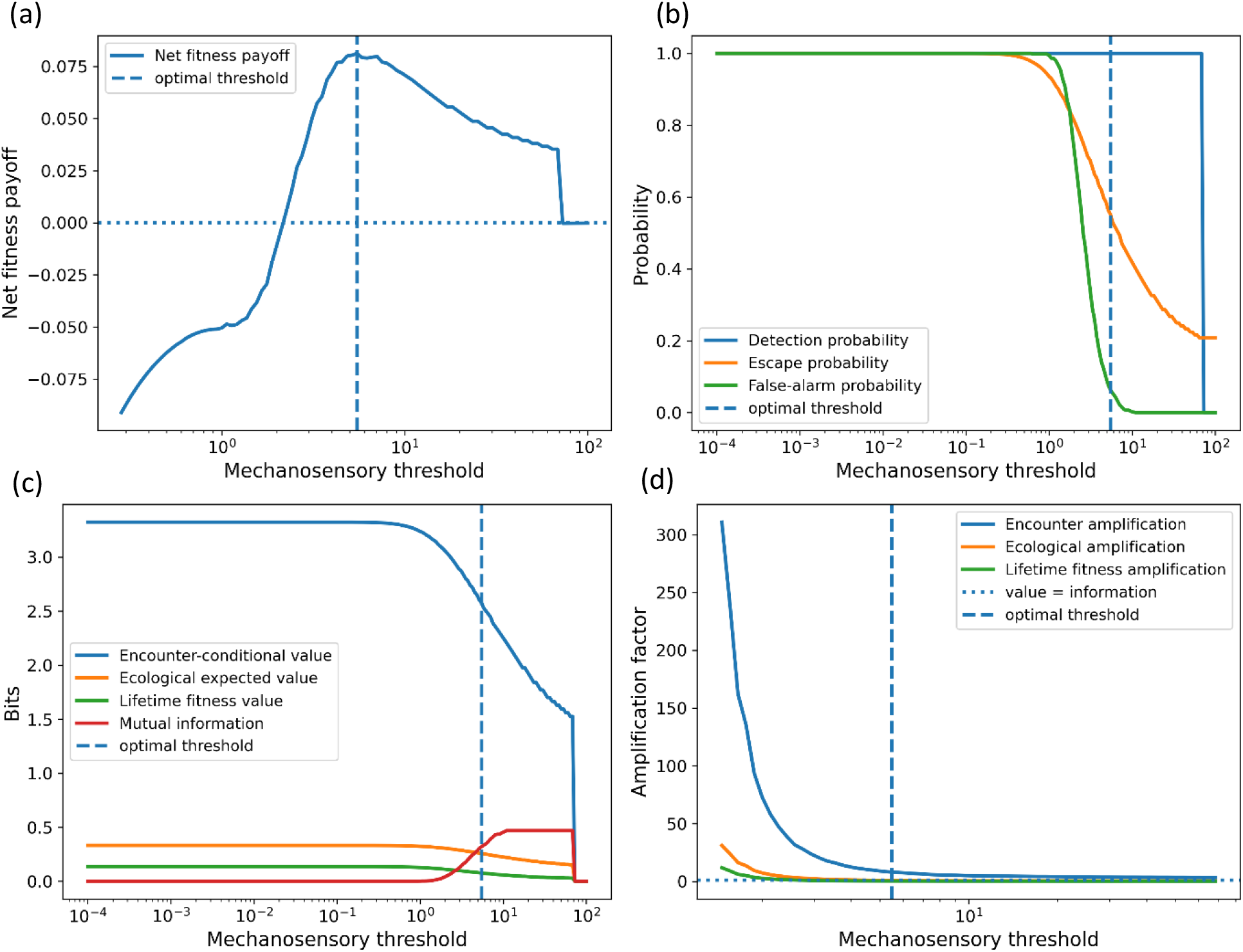
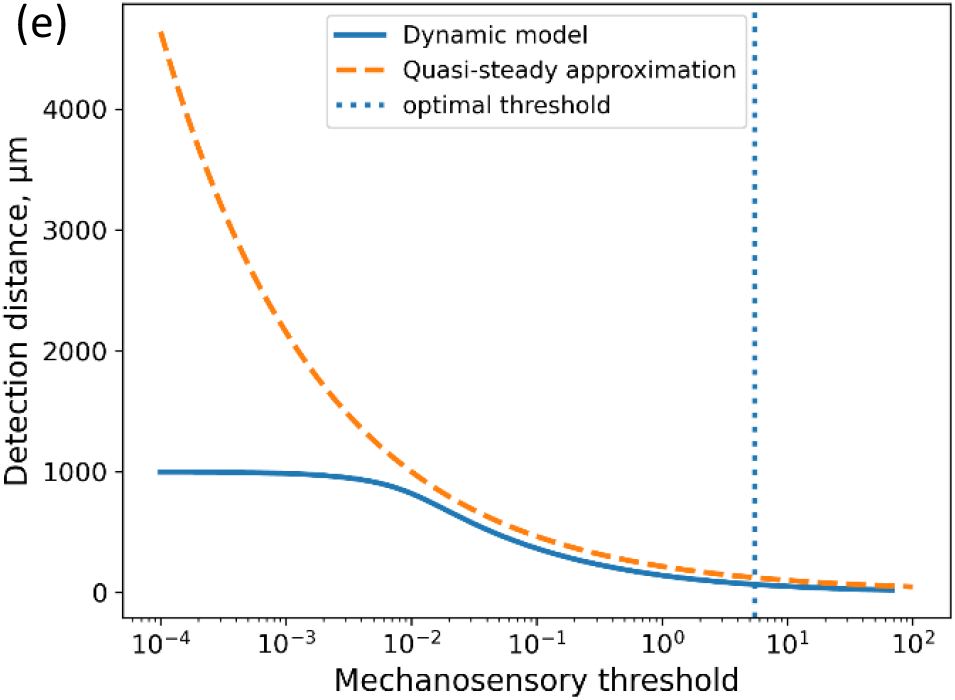
Dependence of prey net fitness payoff, sensory performance, biological value of information, amplification factor, and detection distance on mechanosensory threshold (Θ). (a) Net payoff as a function of mechanosensory threshold, showing a unimodal landscape with an optimal threshold at Θ^∗^ ≈ 5.49. (b) Detection probability, escape probability, and false-alarm probability versus mechanosensory threshold. Increasing the threshold suppresses false alarms but reduces escape probability, while detection probability remains close to unity over most of the threshold range. (c) Encounter-conditional value (V_cond_), ecological expected value (V_ecol_), lifetime fitness value (V_W_), and Shannon mutual information (I) as functions of mechanosensory threshold. The biological value measures exceed Shannon information across most of the threshold range, with the hierarchy V_W_ < V_ecol_ < V_cond_ maintained. (d) Amplification factors (V/I) as functions of mechanosensory threshold. Amplification decreases monotonically with increasing threshold; at the optimal threshold, only encounter-level amplification exceeds unity. (e) Detection distance predicted by the full dynamic-memory model and the quasi-steady approximation. The dynamic model consistently predicts shorter detection distances because finite sensory-memory integration delays detection relative to the quasi-steady limit.

#### Warning time

Once detection occurs, the prey gains a finite interval of time before the predator reaches the capture zone. This interval represents the warning or escape time available for initiating an escape response. Assuming that the predator approaches the prey at a constant speed *U*_*p*_, the warning time is given by

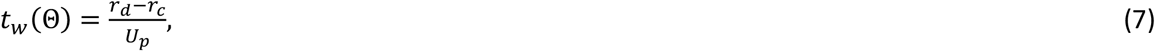

where *r*_*c*_ is the capture distance and *r*_*d*_ is the detection distance. This expression follows directly from predator-prey kinematics and represents the time available for sensory processing, escape initiation, and locomotor response before capture occurs. Because *r*_*d*_ increases with sensory sensitivity (Eq. 6), organisms with lower mechanosensory thresholds obtain longer warning times and consequently greater opportunities to evade predation. The present formulation assumes a direct predator approach at constant swimming speed and therefore provides a simplified description of predator-prey kinematics. In natural systems, predators may accelerate, decelerate, change orientation, or follow more complex trajectories. The constant-speed approximation is adopted here to isolate the relationship between hydrodynamic information, warning time, and survival while retaining analytical tractability.

#### Detection probability

We define a binary detection variable *D* that takes the value 1 when the accumulated sensory evidence reaches the decision threshold before predator capture, and 0 otherwise. Detection therefore occurs when the sensory evidence variable *E*(*t*), governed by Eq. (3), first reaches the threshold value *E* = 1 during predator approach. The threshold *E* = 1 defines the critical evidence level required to trigger a detection event and is chosen as the normalization of the evidence variable. Consequently, *E* < 1 corresponds to insufficient accumulated evidence for reliable predator detection, whereas *E* ≥ 1 indicates that the sensory evidence has reached the decision threshold and an escape response can be initiated.

The detection probability is then defined as

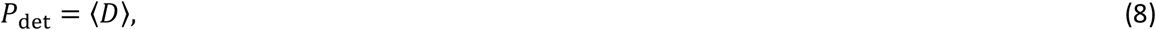

where the angle brackets denote averaging over predator encounters. We retain a threshold rather than a graded detection function because *E* = 1 represents the decision boundary at which accumulated sensory evidence triggers a behavioral response. Thus, detection is binary for an individual deterministic encounter, whereas *P*_det_ becomes graded when averaged over encounters with variation in sensory signals or encounter conditions. A sigmoid mapping could represent additional stochasticity in the sensory decision itself, but would introduce an additional transition-width parameter for which empirical estimates are currently unavailable.

#### Escape success and survival probability

The probability of escaping a predator depends on the warning or escape time (*t*_*w*_) available before capture. We therefore write the escape probability in the general form

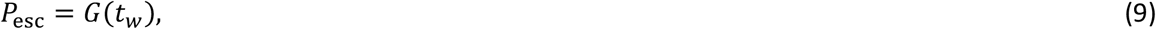

where *G* is a monotonically increasing function. A simple threshold model assumes that escape is successful whenever the warning time exceeds the organism’s reaction time *τ*_*r*_,

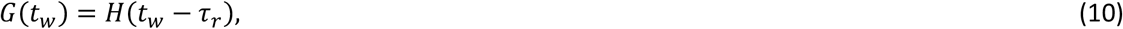

where *H* is the Heaviside step function. This formulation implies that escape probability changes abruptly from 0 to 1 when the available warning time exceeds the reaction time. For numerical calculations and analytical optimization, a smooth approximation is often convenient (Dayan and Abbott 2005),

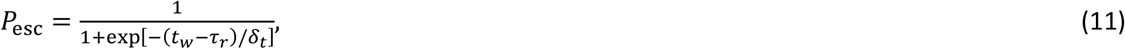

where *δ*_*t*_ characterizes the width of the transition region around the reaction-time threshold. In the limit *δ*_*t*_ → 0, Eq. (11) reduces to the threshold model of Eq. (10). The escape function used here is phenomenological and is intended to capture the central role of warning time in predator avoidance. In real predator-prey interactions, escape success may also depend on prey acceleration, maximum escape speed, turning ability, predator maneuverability, attack angle, body size, and the directionality of the sensory cue. These factors are not modeled explicitly here. Instead, their combined effects are represented by the reaction time *τ*_*r*_ and transition width *δ*_*t*_. Thus, Eq. (11) should be interpreted as a minimal survival-response function rather than a complete biomechanical model of escape behavior.

Survival requires both successful predator detection and successful escape. The overall survival probability during a predator encounter is then

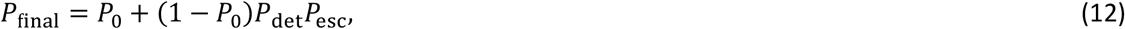

where *P*_0_ represents the baseline probability of survival in the absence of hydrodynamic sensing. The factor (1 − *P*_0_) denotes the fraction of encounters in which the organism would not survive without sensory information. Only this otherwise vulnerable fraction can benefit from hydrodynamic sensing. Thus, the second term represents the additional survival gained when these individuals successfully detect the predator and subsequently escape.

#### False alarms

Hydrodynamic disturbances are not generated exclusively by predators. Turbulence, fluid shear, swimming neighbors, and other background motions can also stimulate mechanoreceptors and potentially trigger escape responses. Such false alarms may incur substantial fitness costs because they interrupt normal activities, waste energy, and reduce opportunities for resource acquisition (Beauchamp and Ruxton 2007). Because predator detection is based on the accumulation of hydrodynamic evidence, false alarms should be defined using the same sensory-memory dynamics as predator detection.

For numerical calculations, the background deformation-rate signal is modeled as a temporally correlated lognormal stochastic process. Specifically,

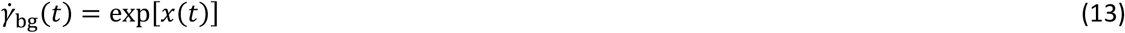

where *x*(*t*) is the logarithm of the background deformation-rate signal and follows a correlated Gaussian process. In discrete time, the logarithmic background signal evolves according to (Gillespie 1996),

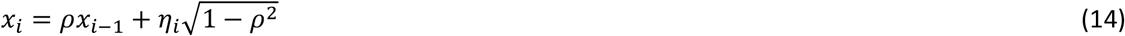

where *η*_*i*_ is a Gaussian random variable,

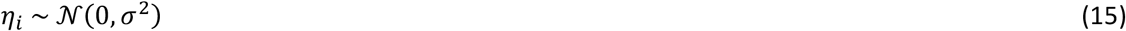

and

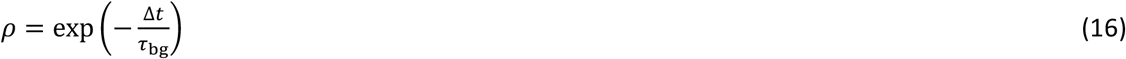

determines the temporal correlation of the background fluctuations. Together, Eqs. (14)–(16) define the exact discrete-time representation of an Ornstein–Uhlenbeck process for the logarithm of the background deformation-rate signal (cf. Eq. (1.10) in (Gillespie 1996)), where *τ*_bg_ is the correlation time and *σ* is the stationary standard deviation of the Gaussian process. Equation (13) then transforms this Gaussian process into a temporally correlated lognormal background deformation-rate field. The same leaky-integrator equation used for predator detection is then applied to this background signal, and *P*_false_ is estimated as the fraction of predator-free observation windows in which *E*_bg_ crosses the detection threshold. This formulation ensures that true detections and false alarms are generated by the same sensory mechanism.

We therefore introduce a background-driven evidence variable *E*_bg_(t), governed by

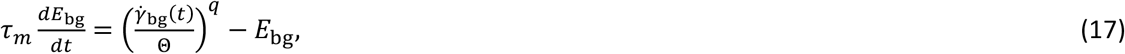

where 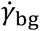 is the background deformation-rate signal experienced in the absence of a predator. A false alarm occurs when background fluctuations alone cause the accumulated evidence to reach the decision threshold,

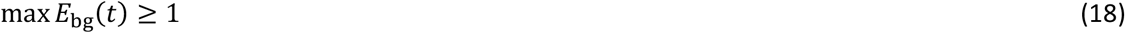

during an observation window *T*. The false-alarm probability is therefore

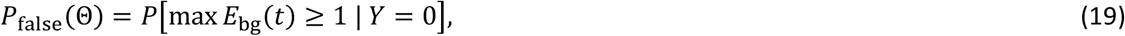

where *Y* = 0 denotes predator absence.

#### Net fitness-payoff consequences of hydrodynamic information

The evolutionary benefit of mechanosensing arises from the increase in survival probability enabled by hydrodynamic information, whereas costs arise from escape behavior, false alarms, and the energetic investment required for the sensory machinery. We therefore define the expected net fitness payoff, *F*(Θ), relative to a no-sensing reference state, associated with a mechanosensory threshold Θ (s^−1^) as:

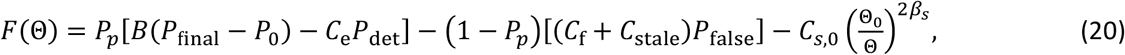

where *P*_*p*_, *P*_final_, *P*_0_, *P*_det_, and *P*_false_ are the predator encounter probability, the survival probability during a predator encounter when hydrodynamic sensing is used, the baseline survival probability in the absence of hydrodynamic sensing, the probability of detecting the predator before capture, and the probability of a false alarm in the absence of a predator, respectively. The coefficients *B, C*_e_, *C*_f_, *C*_stale_, and *C*_*s*,0_ are positive, dimensionless payoff coefficients that express the corresponding biological benefits and costs on a common fitness-payoff scale. Specifically, *B* scales the payoff contribution of an information-induced increase in survival probability; *C*_e_ scales the cost associated with the energetic expenditure and opportunity costs of initiating an escape response following predator detection; *C*_f_ scales the cost of responding to a false alarm; *C*_stale_ scales the additional cost arising when false sensory information persists in memory after the triggering disturbance has disappeared; and *C*_*s*,0_ scales the reference cost of constructing, operating, and maintaining the sensory machinery at Θ = Θ_0_. Although dimensionless, these coefficients are not probabilities or normalized weights and therefore are not restricted to the interval [0, 1]. Consequently, *F*(Θ) is not a probability and is not constrained to the interval [0, 1]. As in other biological cost–benefit models, a net fitness payoff may be negative when the associated costs exceed the benefits (Holland et al. 2002; Both and Visser 2003). Thus, positive values of payoff indicate a net fitness benefit of mechanosensing relative to the no-sensing reference state, whereas negative values indicate that the associated costs exceed the fitness benefit. In the no-sensing limit Θ → ∞, *P*_det_ → 0, *P*_false_ → 0, *P*_final_ → *P*_0_, and the sensory-investment term vanishes, such that *F*(Θ) → 0.

We further assume that achieving greater sensory sensitivity requires a greater investment in effective mechanosensory sensing elements, such that the energetic cost of sensing scales as 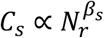, where *N*_*r*_ is the effective number of independent sensing elements while the dimensionless exponent *β*_*s*_ controls how rapidly the sensory cost increases with increasing sensitivity. We assume that the mechanosensory detection threshold decreases with the square root of the number of independent sensing elements, 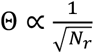. A derivation of this scaling is provided in the Appendix. Combining these relationships yields 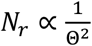, leading to the threshold-dependent sensory cost, 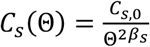, such that increasingly sensitive mechanosensory systems require progressively greater energetic investment in the construction, operation, and maintenance of the sensory machinery. Θ_0_ in Eq. (20) is a fixed reference mechanosensory threshold introduced solely to nondimensionalize the ratio Θ_0_/Θ.

Throughout this model, all benefits and costs are expressed on a common dimensionless fitness-payoff scale. Such a common currency is required because biological benefits and costs measured in different currencies cannot in general be directly combined (Holland et al. 2002). The increase in survival probability, *P*_final_ − *P*_0_, cannot be combined directly with energetic expenditures or opportunity costs merely because all of these quantities can be made dimensionless. Instead, the payoff coefficients act as conversion factors that map each biological consequence onto the same payoff scale. For example, if a sensory cost is measured as a fraction Δ*E*/*E*_budget_ of the organism’s energy budget, its corresponding payoff cost may be written as *λ*_*E*_Δ*E*/*E*_budget_, where *λ*_*E*_ is the marginal contribution of energetic expenditure to the payoff scale. The value of *λ*_*E*_ will generally depend on species, physiological state, resource availability, and the timescale over which the biological consequences are evaluated. Because these conversion relationships are poorly understood for most microbial predator– prey systems, the present model treats the payoff coefficients generically and explores them over broad ranges (see Appendix). Future species-specific studies could derive these payoff coefficients directly from first-principles energetic models and experimentally measured fitness consequences (Laughlin et al. 1998; Beauchamp and Ruxton 2007; Visser 2007; Niven and Laughlin 2008; Meunier et al. 2013; Sartori et al. 2014; Chakraborty et al. 2019; Fahimi et al. 2026*b*, 2026*a*), thereby establishing an explicit quantitative connection between information value, metabolism, and fitness.

#### Evolutionarily optimal threshold

Within the present optimization framework, we define the optimal mechanosensory threshold as the value of Θ that maximizes the expected payoff defined by Eq. (20). The evolutionarily optimal threshold is therefore defined as

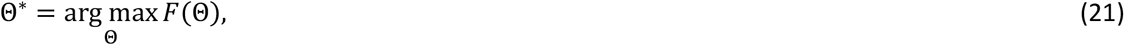

where *F*(Θ) is given by Eq. (20). This threshold represents the optimal trade-off between the benefits of early predator detection and the costs associated with escape behavior, false alarms, stale information, and sensory machinery. Thresholds below Θ^∗^ increase sensitivity but may lead to excessive responses to environmental noise, whereas thresholds above Θ^∗^ reduce false alarms at the expense of delayed predator detection. Consequently, Θ^∗^ identifies the mechanosensory strategy that maximizes the net payoff under a given ecological setting. The payoff-maximizing threshold Θ^∗^ then provides the basis for quantifying the biological value of hydrodynamic information.

#### Mutual information

To quantify the information carried by hydrodynamic signals, we consider each observation window as being in one of two environmental states and represent predator presence by the binary random variable *Y*. Specifically, *Y* = 1 denotes that a predator encounter occurs during the observation window, with probability *p*(*Y* = 1) = *P*_*p*_, whereas *Y* = 0 denotes predator absence, with probability *p*(*Y* = 0) = 1 − *P*_*p*_. The sensory response is similarly represented by the binary random variable *R*, where *R* = 1 indicates detection and *R* = 0 indicates no detection. This representation allows the amount of information conveyed by the sensory system about predator presence to be quantified using Shannon’s mutual information. Similar information-theoretic approaches have been used to quantify how much cells can learn about external signals in bacterial quorum-sensing networks, where multiple environmental signals are integrated through shared signaling pathways (Mehta et al. 2009).

The mutual information between predator presence and sensory response is

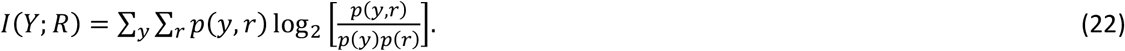

Here, *Y* and *R* denote the environmental-state and sensory-response random variables, respectively, whereas *y* ∈ {0,1} and *r* ∈ {0,1} denote their possible realizations over which the summation is performed. Mutual information measures the reduction in uncertainty about predator presence obtained from the sensory response. It therefore quantifies the informational content of the hydrodynamic cue but does not directly measure its biological utility or fitness consequences. The central objective of the present study is to compare informational content with biological value. This comparison is developed in the following section through a hierarchy of value measures and associated amplification factors.

With predator encounter probability *P*_*p*_, detection probability *P*_det_, and false-alarm probability *P*_false_, the joint probabilities are

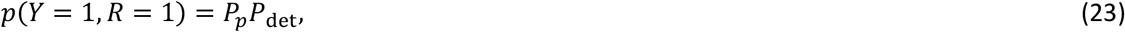

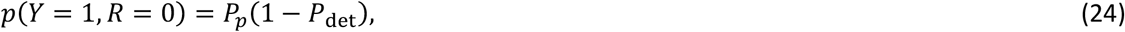

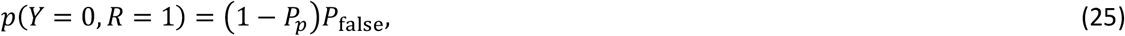

and

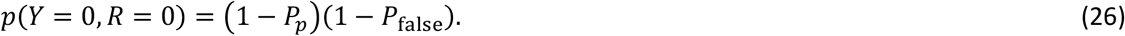

The binary representation adopted here is intentionally chosen to address the biological question considered in the present study, namely whether the prey correctly detects predator presence. Consequently, the mutual information quantifies the Shannon information acquired about predator presence. Ecological and physical variables such as predator size, predator speed, hydrodynamic signal strength, sensory memory, and environmental noise are incorporated implicitly through their effects on the optimal detection threshold Θ^∗^ and the resulting detection (*P*_det_) and false-alarm (*P*_false_) probabilities (see Figure 2). Substitution of Eqs. (23)–(26) into Eq. (22) yields the mutual information used in the numerical calculations.

#### Hierarchical value of information

The preceding formulation quantifies how hydrodynamic information affects survival during predator encounters (Fig. 1). However, the biological value of information depends on the scale at which it is evaluated. A signal may be highly valuable during a predator encounter, yet contribute little to long-term fitness if encounters are rare. We therefore distinguish three complementary measures of information value corresponding to encounter, ecological, and evolutionary scales.

First, we define the encounter-conditional value following Eq. (1)

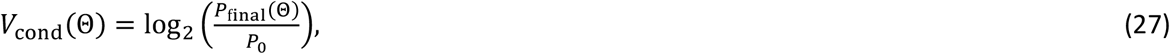

which measures the increase in the probability of surviving a predator encounter attributable to hydrodynamic information. Here, *P*_final_ and *P*_0_ denote the probabilities of surviving a predator encounter with and without mechanosensory information, respectively. Thus, *V*_cond_ answers the question: How valuable is the acquired information, conditional on a predator encounter occurring?

Second, predator encounters occur only with probability *P*_*p*_. Because predator encounter frequencies vary widely among species and environments and depend on factors such as predator and prey densities, swimming speeds, encounter radius, and behavior (Gerritsen and Strickler 1977; Rothschild and Osborn 1988; Almeda et al. 2017), *P*_*p*_ was treated as a free ecological parameter and explored across a broad range of values (see Appendix). This approach allows the model to examine how the value of information depends on encounter rarity independently of any particular predator-prey system. The expected ecological value of information is therefore

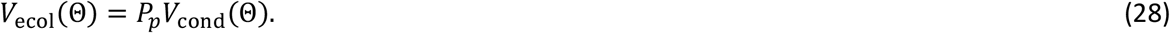

This quantity measures the average value of hydrodynamic information across ecological time by accounting for the fact that predator encounters are intermittent rather than continuous.

Finally, we extend the value-of-information definition in Eq. (1) to survival averaged across predator encounters and non-encounters:

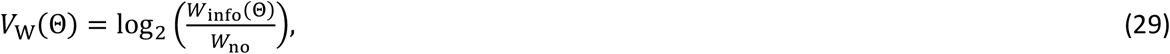

where *W*_info_ and *W*_no_ are the average survival probabilities with and without mechanosensory information, respectively. Because *W*_no_ is constant, a larger *W*_info_(Θ) always corresponds to a larger *V*_W_(Θ). The logarithmic transformation therefore does not change which mechanosensory thresholds perform better or worse, but expresses the proportional survival advantage conferred by information in the same form as the Volkenstein value measure in Eq. (1), facilitating comparison with the other information-value measures and with Shannon information. The present formulation is a mean-field description that averages survival outcomes across predator encounters and non-encounters and does not explicitly incorporate individual heterogeneity, population dynamics, demographic structure, reproduction, or evolutionary-genetic processes.

During a fraction *P*_*p*_ of time, organisms experience predator encounters and survive with probability *P*_final_. During the remaining fraction 1 − *P*_*p*_, no predator is present. The average survival of an information-using organism is therefore

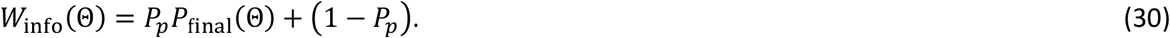

Equation (30) assumes that all non-predation sources of mortality either occur independently of mechanosensory information or affect information-using and non-information-using organisms equally. Consequently, only predator-induced mortality contributes to the difference in expected survival-based fitness quantified here.

In contrast, an organism lacking hydrodynamic sensing cannot benefit from predator detection and therefore survives predator encounters only with probability *P*_0_. Its average survival is therefore

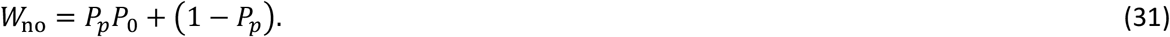

To compare biological value with information content, we normalize each value measure by the mutual information *I*(Θ) between predator presence and sensory response

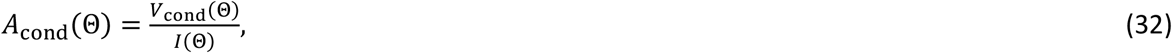

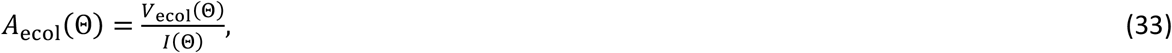

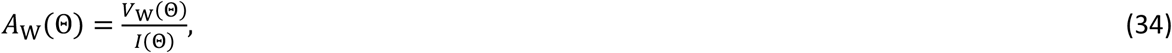

where *I*(Θ) denotes the Shannon mutual information between predator presence and the sensory response as a function of the mechanosensory threshold Θ, as defined in Eq. (22). The amplification factors introduced here may quantify the efficiency with which biological systems convert informational inputs into fitness -relevant consequences. Large amplification factors indicate situations in which relatively small reductions in uncertainty exert disproportionately large effects on survival and evolutionary success. The computational workflow and the order in which the governing equations are implemented are summarized in Figure 2.

## Results and Discussion

Because the theoretical framework developed here is intended to be generally applicable to mechanosensory predator–prey interactions rather than to a specific species, the parameter values listed in Table 1 (Appendix) should be regarded as representative rather than species-specific. These values span broad ranges encompassing ecological scenarios in which benefits either exceed or are outweighed by the various costs associated with escape responses, false alarms, stale information, and sensory investment. The *baseline* coefficients (Table 1 – Appendix) were chosen to represent a biologically plausible regime in which the survival benefit of successful predator detection is large, false alarms impose substantial opportunity costs through unnecessary behavioral responses, reduced resource acquisition, and wasted energy (Beauchamp and Ruxton 2007), and the costs associated with escape responses, stale information, and sensory investment are of comparable magnitude, despite escape responses potentially being energetically expensive in some organisms (e.g., rapid escape jumps and defensive toxin production (Chakraborty et al. 2019; Fahimi et al. 2026*b*)). These values are not species-specific estimates; rather, they define a representative scenario, with robustness assessed by a comprehensive global sensitivity analysis over the full parameter space.

Figure 2 summarizes the computational workflow of the theoretical framework. *Stage I* uses the model equations to identify the evolutionarily favored mechanosensory threshold, Θ^∗^, by evaluating the net fitness-payoff function over a range of candidate thresholds. *Stage II* then uses this optimal threshold to quantify the mutual information, the encounter-conditional, ecological, and lifetime fitness values of information, and the corresponding amplification factors. This workflow highlights the logical progression from mechanosensory signal detection and evolutionary optimization to the biological evaluation of information.

Optimization of the mechanosensory threshold revealed a strong separation between information content and biological value. The evolutionarily optimal threshold was Θ^∗^ ≈ 5.49, corresponding to a mutual information of only 0.32 bits between predator presence and sensory response. Despite this modest information content, the encounter-conditional value of information reached 2.58 bits, yielding an amplification factor of 8.04. Thus, a relatively small amount of sensory information can generate a disproportionately large survival benefit when used to guide predator-avoidance decisions.

At the optimal threshold, predator detection was reliable (*P*_det_ = 1.0), escape success remained substantial (*P*_esc_ ≈ 0.55), and false alarms were infrequent (*P*_false_ ≈ 0.06). For the representative parameter values used here, the resulting dynamic detection distance was approximately 66 μm, corresponding to a warning time of approximately 0.05 s before predator capture. These values depend on the assumed predator and prey characteristics, swimming velocities, and sensory parameters, and are presented as an illustrative example rather than universal predictions. These values emerged from a trade-off between sensitivity and reliability: lower thresholds increased false alarms, whereas higher thresholds reduced warning time and escape success. The optimum therefore represents the best compromise between acquiring information about predators and avoiding the costs of unnecessary responses.

The hierarchical value framework reveals that information contributes differently across biological scales. While the encounter-conditional value was large (*V*_cond_ = 2.58 bits), ecological averaging reduced the expected value to 0.26 bits, and the lifetime fitness value was further reduced to 0.08 bits. This decline reflects the fact that predator encounters constitute only a fraction of ecological time, so the benefits of information are diluted when averaged across both predator and non-predator conditions. Nevertheless, the optimal sensory system increased the survival-based fitness measure from *W*_no_ = 0.91 to *W*_info_ = 0.96, corresponding to a relative survival advantage of approximately 5.5% (i.e. [(0.96 – 0.91)/0.91] × 100). Consequently, the optimized sensory strategy produces a measurable survival advantage relative to the absence of mechanosensory information. This comparison quantifies the overall benefit of information use, whereas selection on the mechanosensory phenotype itself depends on how the underlying payoff changes continuously with the sensory threshold Θ, examined below.

To evaluate robustness, we repeated the stochastic false-alarm calculation across 20 independent random seeds and performed a global sensitivity analysis over 120 parameter combinations. The random-seed analysis demonstrated stable numerical behavior despite stochastic variation in the background-noise realizations. Across the 20 seeds, Θ^∗^ = 5.78 ± 0.51, *V*_cond_ = 2.55 ± 0.05, *V*_ecol_ = 0.25 ± 0.005, *V*_W_ = 0.075 ± 0.003, and *A*_cond_ = 7.63 ± 0.72, *A*_ecol_ = 0.76 ± 0.07, *A*_W_ = 0.22 ± 0.02 (mean ± SD). In the global sensitivity analysis, *V*_W_ remained positive in 86% of parameter combinations, the value hierarchy held in 86%, encounter amplification occurred in 80%, ecological amplification occurred in 21%, and lifetime fitness amplification occurred in 7%. Across the sensitivity ensemble, the median encounter-conditional value was 2.00 bits despite a median mutual information of only 0.41 bits, indicating that substantial biological value can arise from very small amounts of sensory information. These results show that value amplification is common at the encounter scale but becomes increasingly conditional at the ecological and lifetime scales. Figures 3a–e and 4a–c present a representative example selected from the 120 parameter combinations examined in the global sensitivity analysis.

The optimization framework adopted here identifies the mechanosensory threshold that maximizes the net payoff under the assumed ecological conditions. It does not, however, model the evolutionary dynamics or the distribution of phenotypes within a population. The realized population mean need not coincide with the payoff-maximizing threshold, particularly when the payoff landscape is asymmetric in the presence of phenotypic variation or when the phenotypic distribution itself is skewed (Urban et al. 2013), while mutation, genetic architecture, and random genetic drift can further cause evolved mean phenotypes to deviate from selective optima (Lynch 2012, 2020; Lynch and Menor 2025). We therefore interpret Θ* as the payoff-maximizing threshold under the assumptions of the present model, rather than as a predicted evolutionary endpoint.

At low mechanosensory thresholds (high sensitivity), the net fitness payoff relative to the no-sensing reference state is negative because the combined costs outweigh the survival benefits (Fig. 3a). As the threshold increases, net payoff increases rapidly and reaches a maximum at the optimal threshold (Θ* ≈ 5.49). Beyond this optimum, net payoff declines progressively with further increases in mechanosensory threshold. The resulting unimodal payoff landscape indicates that neither very low nor very high thresholds maximize net payoff; instead, an intermediate threshold yields the highest net payoff and represents the optimal mechanosensory threshold. Because Θ is the focal mechanosensory trait in the present model, the local effect of changes in this trait is given by the slope *dF*/*d*Θ of the payoff landscape. The gradient is positive below the payoff-maximizing threshold and negative above it, with *dF*/*d*Θ = 0 at Θ*.

Figure 3b shows how detection probability, escape probability, and false-alarm probability vary with mechanosensory threshold. Detection probability remains close to unity over a broad range of thresholds and decreases only at very high thresholds. In contrast, escape probability declines progressively as the mechanosensory threshold increases, reflecting the shorter warning times associated with later detection. False-alarm probability exhibits the opposite trend, remaining high at low thresholds but falling rapidly as the threshold increases and approaching zero at higher thresholds. At the optimal threshold (Θ* ≈ 5.49), detection probability is still effectively maximal, escape probability remains moderate, and false-alarm probability has been substantially reduced. These opposing trends demonstrate that increasing the mechanosensory threshold suppresses false alarms at the expense of escape performance while leaving detection probability largely unaffected over most of the threshold range.

The three biological value measures exhibit distinct dependencies on mechanosensory threshold (Fig. 3c). Across most of the threshold range, the encounter-conditional value is the largest measure, followed by the ecological expected value and the lifetime fitness value. All three measures remain nearly constant at low thresholds and decline progressively as the mechanosensory threshold increases. At the optimal threshold (Θ* ≈ 5.49), the hierarchy *V*_W_ < *V*_ecol_ < *V*_cond_ is clearly maintained. In contrast, Shannon mutual information is close to zero at low thresholds (because at low thresholds, frequent responses to both predators and background noise provide little information about predator presence), increases with mechanosensory threshold to a maximum near the optimal threshold, and remains substantially smaller than the encounter-conditional value throughout most of the threshold range. These results demonstrate that the biological value of sensory information and its Shannon information content exhibit markedly different dependencies on mechanosensory threshold.

Figure 3d compares the amplification factors obtained by dividing each biological value measure by the corresponding Shannon mutual information. All three amplification factors decrease monotonically with increasing mechanosensory threshold, indicating that amplification is greatest at low thresholds and progressively weakens as the threshold increases. The encounter amplification factor remains the largest across the entire threshold range, followed by the ecological and lifetime fitness amplification factors. At the optimal threshold (Θ* ≈ 5.49), only the encounter amplification factor exceeds unity, whereas the ecological and lifetime fitness amplification factors remain below unity. The amplification is strongest for encounter-conditional value and weakest for lifetime fitness value, with the latter approaching unity most closely.

The full dynamic model includes finite sensory-memory dynamics, whereas the quasi-steady approximation assumes instantaneous equilibration of the internal sensory state. The full dynamic-memory model predicts consistently shorter detection distances than the quasi-steady approximation across the entire range of mechanosensory thresholds (Fig. 3e). Consequently, predators must approach more closely before detection occurs in the dynamic model than in the quasi-steady approximation. The discrepancy is greatest at low thresholds and progressively decreases with increasing threshold, where both models predict relatively short detection distances. At the optimal threshold (Θ* ≈ 5.49), the dynamic model predicts a detection distance of approximately 66 μm, compared with approximately 122 μm for the quasi-steady approximation. This difference reflects the finite accumulation time required for sensory evidence in the dynamic model, which delays detection relative to the quasi-steady approximation.

The ecological amplification map (Fig. 4a) shows a transition from negative to positive log_10_ *A*_ecol_, corresponding to a transition from sub-unity to greater-than-unity amplification. This transition occurs across the full range of false-alarm costs, although the magnitude of amplification varies with false-alarm cost. Consequently, information that is highly valuable during an encounter contributes progressively more to ecological value as predator encounters become more frequent.

**Figure 4:**
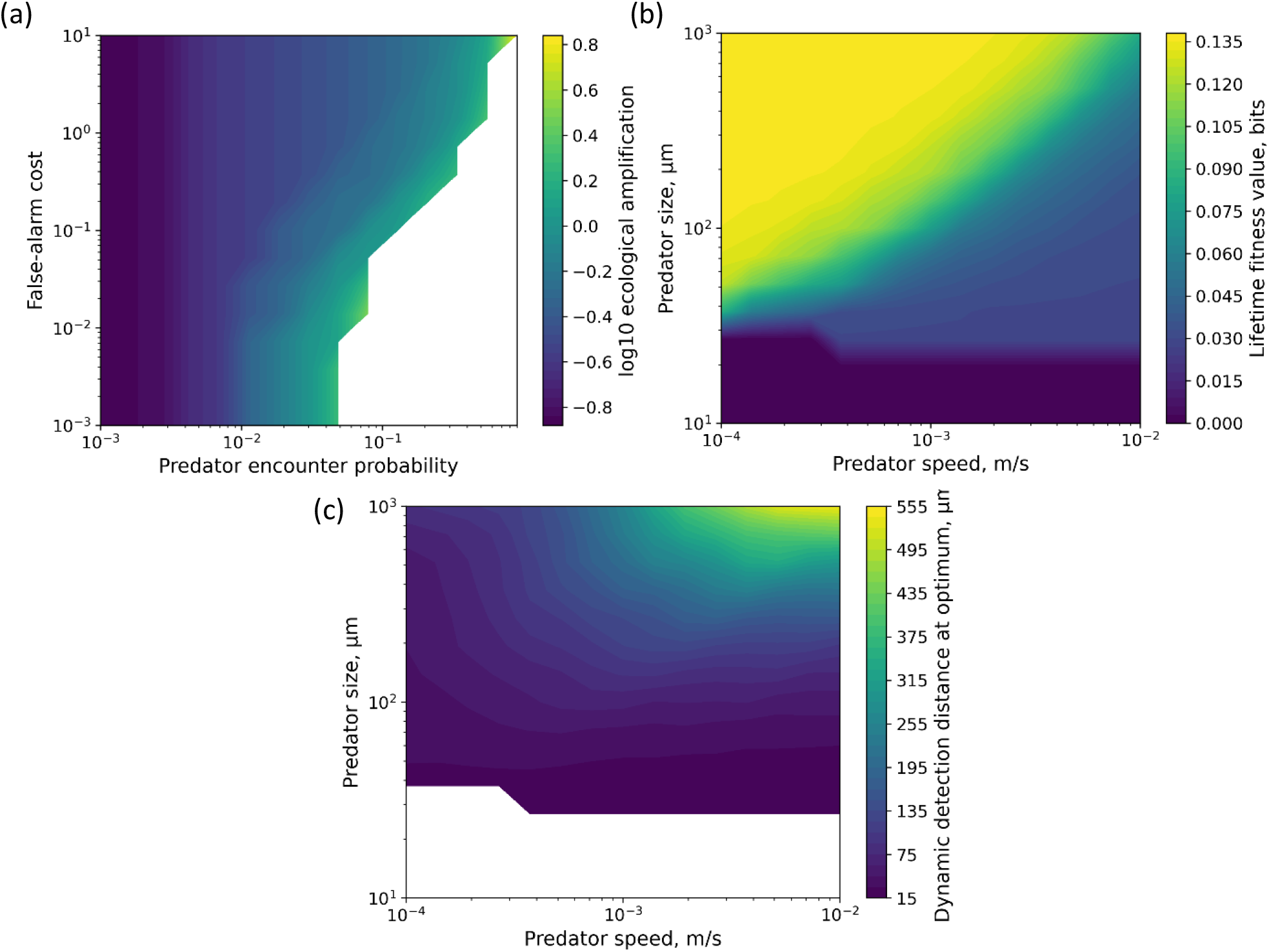
Ecological amplification and lifetime fitness value of mechanosensory information. (a) Ecological amplification as a function of predator encounter probability and false-alarm cost, showing a transition from negative to positive log_10_ A_ecol_, or equivalently from sub-unity to greater-than-unity amplification, as predator encounters become more frequent. (b) Lifetime fitness value of mechanosensory information for the prey as a function of predator size and swimming speed, with the greatest value occurring for large, slow-moving predators that generate strong hydrodynamic signals while providing sufficient warning time for escape. (c) Corresponding hydrodynamic detection distance as a function of predator size and swimming speed. Detection distance increases with both predator size and speed because stronger hydrodynamic deformation rates exceed the mechanosensory threshold at greater distances. Although faster predators are detected earlier, their higher approach speeds shorten the available warning time, explaining why increased detection distance does not necessarily produce higher lifetime fitness value.

The lifetime fitness value of mechanosensory information for the prey depends strongly on predator size and swimming speed (Fig. 4b). Within the present model, this value generally increases with predator size and decreases with predator swimming speed. The highest values occur when predators are large and move slowly, whereas the lowest values occur when predators are small across much of the speed range. These trends arise because larger predators generate stronger hydrodynamic disturbances and slower predators provide more time for an effective escape response. This analysis isolates the consequences of predator size and speed for detection by the prey; it does not account for possible size-dependent differences in the predator’s ability to detect, pursue, or capture the prey, or in the prey’s ability to escape after detection. Such effects could modify the relationship between predator size and the value of mechanosensory information. Because Fig. 4b represents a single illustrative parameter set, the quantitative values also depend on the assumed predator encounter probability (*P*_*p*_), which may vary with predator abundance and was explored in the global sensitivity analysis.

The hydrodynamic basis of the patterns observed in Fig. 4b is illustrated by the corresponding detection-distance map in Fig. 4c. Detection distance increases with both predator size and swimming speed because larger and faster predators generate stronger hydrodynamic deformation rates that exceed the mechanosensory threshold at greater distances. Detection distances exceeding several hundred micrometres occur for the largest and fastest predators considered, whereas small predators are detected only at relatively short ranges. Although faster predators are detected farther away, they also reduce the available warning time by closing the remaining distance more rapidly, explaining why increased detection distance does not necessarily translate into greater lifetime fitness value in Fig. 4b.

### BOX 3. Limitations of the Study

The hydrodynamic model employs a far-field stresslet approximation for predator-generated flows. Although this captures the dominant scaling of hydrodynamic disturbance strength, real microorganisms may experience more complex near-field flow structures that depend on body geometry, propulsion mechanism, orientation, and fluid boundary conditions. Incorporating organism-specific flow fields would improve quantitative predictions of detection distance and warning time.

The baseline model treats predator approach, sensory parameters, and hydrodynamic geometry deterministically. As a result, detection is represented as a threshold-crossing outcome rather than a fully stochastic probability. Future extensions should incorporate variation in predator trajectories, accelerations, turning behaviors, heterogeneous encounter geometries, approach angles, sensory noise, and receptor variability to estimate genuinely probabilistic detection functions.

Escape success is modeled as a simple function of warning time. This formulation does not explicitly include prey locomotor mechanics, predator maneuverability, directionality of the escape response, or attack geometry. More detailed biomechanical escape models could be incorporated in future extensions, but the present formulation is sufficient for isolating how hydrodynamic information changes survival probability.

The net fitness-payoff function employed here is intentionally generic rather than species-specific. Although it captures the major trade-offs associated with predator detection, including the benefits of enhanced survival and the costs of escape responses, false alarms, and sensory investment, alternative formulations could yield quantitatively different optimal thresholds and amplification factors. In particular, the present framework does not explicitly account for the effects of mechanosensory behavior on resource acquisition, growth, or reproductive output, which may further influence the evolutionarily favored mechanosensory threshold. Future species-specific models could incorporate these additional life-history components within the general framework developed here.

The present framework optimizes only the mechanosensory detection threshold while treating the sensory architecture itself as fixed. In reality, natural selection may also optimize the number, density, spatial distribution, and localization of mechanoreceptors, balancing improvements in sensing accuracy against the energetic costs of constructing and maintaining sensory systems (Berg and Purcell 1977; Lynch 2024). Extending the present framework to include the evolution of sensory architecture would provide a closer connection to theories of optimal chemosensing and environmental sensing.

The present framework is sufficiently general to describe mechanosensory predator detection in both unicellular plankton and multicellular zooplankton. However, the biological interpretation and parameterization of the sensory components differ among taxa. In unicellular organisms, sensing is generally mediated by mechanosensitive ion channels responding to fluid deformation (Árnadóttir and Chalfie 2010), whereas in zooplankton it typically involves specialized mechanoreceptive setae coupled to the nervous system (Shen et al. 2020). Future applications should therefore employ taxon-specific sensory architectures and parameter values.

The mutual-information analysis is based on binary environmental and sensory states. Real hydrodynamic signals contain continuous information regarding signal magnitude, temporal dynamics, predator trajectory, and predator identity. Incorporating continuous-state information measures would provide a more complete description of sensory information processing and may alter quantitative amplification factors while preserving the conceptual distinction between information content and biological value.

The present framework does not explicitly incorporate adaptive learning or experience-dependent updating of sensory decision-making during repeated predator encounters (Kikuchi and Sherratt 2015; Sherratt and Peet-Paré 2017; Gershman et al. 2021; Gunawardena 2022; Heerwig et al. 2023; Eckert et al. 2024). Such processes could alter the frequency of decision errors, including missed detections and false alarms, by allowing organisms to adjust sensory thresholds based on past experience. Future work could therefore examine how the balance between genetically encoded sensory strategies and experience-dependent learning shapes information processing and the biological value of information over time.

The present framework assumes that predator encounters occur independently with probability *P*_*p*_ and therefore neglects temporal clustering, spatial heterogeneity, and predator aggregation. In natural environments, encounters may be correlated through patchy predator distributions, seasonal fluctuations, or repeated interactions with the same predator population. Such correlations could alter the ecological and evolutionary value of information by changing the temporal distribution of predation risk.

## Conclusion

This study develops a quantitative framework for evaluating the biological value of environmental information in microbial predator–prey interactions. By extending Volkenstein’s concept of information value from molecular biology to ecological sensing, the framework distinguishes the amount of information carried by an environmental cue from the biological consequences of using that information. The principal theoretical contribution is the introduction of a hierarchical description of biological information value at three organizational scales: encounter-conditional value, ecological value, and lifetime fitness value. This hierarchy provides a quantitative bridge between Shannon information theory and the ecological and evolutionary consequences of environmental sensing.

The model demonstrates that environmental information can produce substantial biological benefits even when its Shannon information content is modest. The optimal mechanosensory threshold emerges from a trade-off between the benefits of early predator detection and the costs associated with false alarms, escape responses, stale information, and sensory investment. At this optimum, predator detection remains highly reliable while false alarms are substantially reduced and sufficient warning time is retained for successful escape. These results illustrate that natural selection may favor sensory strategies that maximize biological value rather than simply maximizing the amount of information acquired.

The hierarchical analysis further shows that the biological value of environmental information is inherently scale-dependent. During predator encounters, relatively small reductions in uncertainty can generate disproportionately large improvements in survival, producing encounter-level value amplification. However, this value becomes progressively diluted when averaged across ecological time and lifetime survival because predator encounters are intermittent and information use incurs biological costs. Consequently, the same sensory cue can possess markedly different biological values depending on the organizational scale at which it is evaluated.

The global sensitivity analysis further demonstrates that this hierarchical framework is robust across a broad range of biologically plausible parameter combinations. Although the magnitude of information value and amplification depends on ecological conditions and sensory costs, the qualitative distinction between Shannon information and biological value persists over much of the explored parameter space. Value amplification is therefore common at the encounter scale but becomes increasingly conditional as broader ecological and lifetime processes are incorporated.

Although the present study focuses on hydrodynamic predator avoidance in planktonic microorganisms, the underlying framework is considerably more general. Any biological system in which organisms acquire environmental information to make threshold-based decisions—including chemosensing, visual sensing, mechanosensing, prey detection, nutrient acquisition, and toxin avoidance—can, in principle, be analyzed using the same approach. Rather than simply asking whether information is beneficial, the framework links measurable physical, sensory, and ecological variables to the biological consequences of information use. Given estimates of signal strength, sensory threshold and memory, environmental noise, encounter frequency, and the costs of sensing and responding, the model can be used to identify the threshold that maximizes net fitness payoff, quantify the information acquired, and determine how its biological value changes from individual encounters to broader ecological scales. It can therefore be used to compare sensory strategies among organisms or environments, identify which parameters most strongly limit the value of information, and guide experiments that manipulate signal strength, noise, encounter frequency, or sensory investment.

Ultimately, the framework provides a quantitative toolkit for integrating environmental information into ecological and evolutionary research. The equations developed here can be combined with experimentally measured sensory performance, encounter frequencies, behavioral responses, and fitness consequences to estimate the biological value of information under different ecological conditions. This enables quantitative comparisons among species, sensory modalities, environments, and evolutionary strategies, while providing a common framework for incorporating biologically meaningful information into theoretical models of behavior, ecology, and evolution. We anticipate that this framework will facilitate experimental tests of how natural selection shapes sensory systems and will help establish biological information value as a measurable and predictive quantity alongside information content.

## Acknowledgment

PF gratefully acknowledges the unwavering support of his family. ML is grateful to the National Institutes of Health (2R35GM122566), the National Science Foundation (DBI-2119963, DEB-1927159), the Simons Foundation (735927), and the Moore Foundation (12186).

## Appendix

**Table S1:** Model parameters and numerical settings. Parameter values used for the representative dynamic-memory simulations and parameter scans. Parameters were sampled according to distributions reflecting their expected biological variability. Positive quantities that naturally vary over orders of magnitude (e.g., body size, time constants, and cost coefficients) were sampled from log-uniform distributions so that each order of magnitude was represented equally. Parameters bounded within a relatively narrow interval without evidence for scale-invariant variation (e.g., probabilities, dimensionless exponents, and geometric factors) were sampled from uniform distributions.

| Symbol | Baseline value | Sensitivity range | Distribution | Units | Definition |
| --- | --- | --- | --- | --- | --- |
| $A$ | 1.0 | 0.5 – 2.0 | Log-uniform | --- | Hydrodynamic geometric coefficient in Eq. (2) |
| $a_p$ | $10^{-4}$ | $2 \times 10^{-5} - 5 \times 10^{-4}$ | Log-uniform | m | Predator characteristic radius |
| $U_p$ | --- | --- | --- | $\text{m s}^{-1}$ | Predator swimming speed<br>$v[\mu\text{m. s}^{-1}] = 37(V[\mu\text{m}^3])^{0.13 \pm 0.03}$<br>Reference: (Schavemaker and Lynch 2022) |
| $r_c$ | $2.0 \times 10^{-5}$ | $0.1 a_p - 0.5 a_p$ | Uniform | m | Capture distance |
| $r_0$ | $50 r_c$ | $20 r_c - 100 r_c$ | Uniform | m | Initial predator-prey separation |
| $P_0$ | 0.1 | 0.02 – 0.4 | Uniform | --- | Baseline survival probability without mechanosensory information |
| $P_p$ | 0.1 | 0.01 – 0.3 | Uniform | --- | Predator encounter probability |
| $q$ | 1.0 | 0.7 – 1.5 | Uniform | --- | Sensory nonlinearity exponent |
| $\tau_m$ | 0.15 | 0.03 – 0.5 | Log-uniform | s | Sensory memory time constant |
| $\tau_r$ | 0.04 | 0.01 – 0.1 | Log-uniform | s | Escape reaction time |
| $\delta t$ | 0.03 | 0.01 – 0.07 | Log-uniform | s | Width of escape-response transition |
| $B$ | 2.0 | 0.5 – 5.0 | Log-uniform | --- | Survival-benefit payoff coefficient |
| $C_e$ | 0.02 | 0.01 – 0.5 | Log-uniform | --- | Escape cost coefficient |
| $C_f$ | 0.20 | 0.02 – 1.0 | Log-uniform | --- | False-alarm cost coefficient |
| $C_{\text{stale}}$ | 0.02 | $5 \times 10^{-3} - 0.5$ | Log-uniform | --- | Stale information cost coefficient |
| $C_{s,0}$ | 0.02 | $5 \times 10^{-3} - 0.5$ | Log-uniform | --- | Baseline sensory cost coefficient |
| $\beta_s$ | 0.5 | 0.2 – 1.0 | Uniform | --- | Sensory cost exponent |
| $\sigma_{\text{bg}}$ | 1.0 | 0.3 – 2.0 | Log-uniform | --- | Log-space standard deviation of background deformation |
| $\tau_{\text{bg}}$ | 0.05 | 0.01 – 0.3 | Log-uniform | s | Correlation time of background fluctuations |
| $I_{\text{min}}$ | 0.01 | --- | --- | bits | Minimum mutual information used when computing amplification ratios |
| $T_{\text{obs}}$ | 1.0 | --- | --- | s | Observation interval used to estimate false alarms |
| $\theta$ | --- | $10^{-4} - 10^2$ | --- | $\text{s}^{-1}$ | Mechanosensory threshold search range |
| $\theta_0$ | 1.0 | --- | --- | $\text{s}^{-1}$ | Reference mechanosensory threshold |

### Scaling of the mechanosensory detection threshold with receptor number

Suppose each of the *N*_*r*_ mechanosensory receptors measures the local hydrodynamic disturbance with an independent random measurement error having standard deviation *σ*. When the sensory system combines information from all receptors, the uncertainty of the combined measurement is reduced. Specifically, for *N*_*r*_ independent measurements, the standard deviation of the combined estimate is

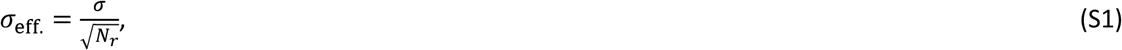

a classical result in statistics known as the standard error of the mean (or, more generally, of independent measurements). Because a hydrodynamic disturbance must exceed the measurement uncertainty to be reliably detected, the minimum detectable disturbance (the detection threshold) is proportional to this effective uncertainty. Therefore,

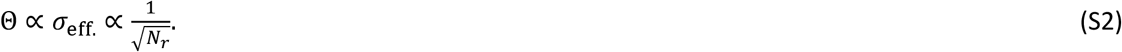

Thus, increasing the number of independent mechanosensory receptors improves detection sensitivity, causing the detection threshold to decrease with the square root of receptor number.

